# An optogenetic tool to raise intracellular pH in single cells and drive localized membrane dynamics

**DOI:** 10.1101/2021.03.09.434608

**Authors:** Caitlin E. T. Donahue, Michael D. Siroky, Katharine A. White

## Abstract

Intracellular pH (pHi) dynamics are critical for regulating normal cell physiology. For example, transient increases in pHi (7.2-7.6) regulate cell behaviors like cell polarization, actin cytoskeleton remodeling, and cell migration. Most studies on pH-dependent cell behaviors have been performed at the population level and use non-specific methods to manipulate pHi. The lack of tools to specifically manipulate pHi at the single-cell level has hindered investigation of the role of pHi dynamics in driving single cell behaviors. In this work, we show that Archaerhodopsin (ArchT), a light-driven outward proton pump, can be used to elicit robust and physiological pHi increases over the minutes timescale. We show that activation of ArchT is repeatable, enabling the maintenance of high pHi in single cells for approximately 45 minutes. We apply this spatiotemporal pHi manipulation tool to determine whether increased pHi is a sufficient driver of membrane ruffling in single cells. Using the ArchT tool, we show that increased pHi in single cells can drive localized membrane ruffling responses within seconds and increased membrane dynamics (both protrusion and retraction events) compared to unstimulated ArchT cells as well as control cells. Overall, this tool allows us to directly investigate the relationship between increased pHi and cell behaviors such as membrane ruffling. This tool will be transformative in facilitating experiments required to determine if increased pHi drives these cell behaviors at the single-cell level.

## INTRODUCTION

In normal epithelial cells, intracellular pH (pHi) is regulated between 7.0-7.2, but transient increases in pHi (7.2-7.6) are linked to cell behaviors including cell polarization^1,2^, cytoskeleton remodeling^3,4^, and directed cell migration.^1,2,5,6^ However, most studies on pH-dependent cellular behaviors manipulate pHi using non-specific methods^7^, which can confound interpretation of biological cause and effect. The lack of appropriate tools to specifically and spatiotemporally manipulate pHi in living cells has obscured our understanding of the molecular mechanisms driving pHi-regulated cell behaviors. Elucidating these molecular mechanisms will advance our understanding of normal pH-regulated behaviors and enable us to identify how dysregulated pHi drives disease states such as cancer (constitutively increased pHi)^7,8^ and neurodegenerative diseases (constitutively decreased pHi)^9,10^.

Current methods to manipulate intracellular pHi are nonspecific and technically challenging.^7^ For example, genetic overexpression^11^ or ablation^12^ of ion transporters alter pHi but lack specificity, as ion transporters can be linked to non-proton gradient changes^13^ and function as signaling scaffolds.^14^ Furthermore, the use of ion transport inhibitors is very common^8,15^ but recent studies have reported significant off-target effects of various pHi-lowering drugs.^16,17^ Micropipette pHi manipulation techniques are highly specific and adaptable to single-cell analysis.^18^ However, micropipette pHi manipulation is technically challenging, has poor temporal control, is damaging to the cell, and is low-throughput.^19^ Finally, while proton uncaging molecules can be photolysed to release protons with good temporal control, spatial control is poor. Also, proton uncaging requires UV photolysis which is cytotoxic and can only be used to lower pHi, limiting experimental applications.^20–23^ The ideal tool to spatiotemporally manipulate pHi in single cells would be non-damaging to the cell, have good spatial and temporal control, be robust and reversible, and allow for repeated pHi manipulations.

We identified Archaerhodopsin (ArchT), an outward proton pump activated by 561 nm light^24–27^, as a potential tool to spatiotemporally raise pHi in single cells. Archaerhodopin is natively light-activatable, expressed on the plasma membrane, and is a viable first-generation pHi manipulation tool. Archaerhodopsins have been previously developed as light-activated neuronal silencers.^24,25^ Under long (15 sec) photoactivation conditions, ArchT did not significantly increase pHi in neurons.^25^ However, some recent work suggests that ArchT could be a viable tool for pHi manipulation. First, a study using an adapted ArchT with ER export for improved surface expression showed increased pHi locally in the spatially-restricted synaptic bouton under very long (2 minute) photoactivation conditions.^26^ Second, ArchT was recently used as a high-resolution spatiotemporal tool to generate proton fluxes and measure gap junction connectivity in mammalian cells and the developing *Drosophila* brain.^27^ Based on these foundational studies, we hypothesized that ArchT could be adapted to reversibly and robustly manipulate intracellular pHi in mammalian cells on the minutes timescale.

In this work, we show the optimization and characterization of ArchT to spatiotemporally raise pHi in single cells. Using optimized photoactivation protocols, we can robustly raise pHi in single cells over the minutes-timescale. The ArchT-induced pHi response can be seen with a range of light powers, and we can induce a range of physiological increases in pHi (0.10-0.47), similar to those reported during pH-regulated processes (0.1-0.35). Furthermore, we show that this tool can be repeatedly photoactivated, allowing for sustained pHi manipulation on a longer timescale (~45 minutes). When we raise pHi spatiotemporally using ArchT in single living cells, we found that increased pHi drives localized membrane protrusion and ruffling responses in single cells. ArchT is a robust optogenetic tool for spatiotemporally increasing cytosolic pH within single-cells.

## METHODS

### Cell Culture

NIH-3T3 mouse embryo fibroblast (NIH-3T3 ATCC^®^ CRL-1658) cells were cultured in Dulbecco’s Modified Eagle’s Medium (DMEM, Corning 10-013-CV) supplemented with 10% Fetal Bovine Serum (FBS, Peak Serum, PS-FB2). Retinal Pigment Epithelial (RPE, ATCC^®^ CRL-4000) cells were cultured in Roswell Park Memorial Institute 1640 media (RPMI-1640, Corning, 10-040-CV), supplemented with 10% FBS (Peak Serum, PS-FB2) and GlutaMAX™ (Gibco, 35050-061) up to 446 mg/L of glutamine. All cells were grown in humidified incubators at 37°C, 5% CO_2_.

### Plasmid Constructs

pcDNA3.1-ArchT-BFP2-TSERex was a gift from Yulong Li (Addgene plasmid # 123312) pCDNA3-mCherry-SEpHluorin (mCherry-pHluorin) was a gift from Sergio Grinstein (Addgene plasmid #32001)

### Transfection Protocol

RPE and NIH-3T3 cells were transiently transfected using Lipofectamine 2000 (Life Technologies, 11668-019) Briefly, a 1:1 ratio of DNA-Lipofectamine complex was prepared in 1 mL serum-free media (DNA: 2 μg of a single-construct or 1μg of each construct for double transfection) (Lipofectamine used at 3:1 ratio to DNA) and incubated at room temperature for 5 minutes. This was then combined and added to 1 mL serum-free media on cells. Cells were incubated with transfection media for 8 hours at 37°C with 5% CO_2_, before exchanging with complete media. Cells were imaged the following day.

### Microscopy

All imaging experiments were performed using a Nikon Ti2 Eclipse confocal microscope equipped with a spinning disk (CSU-X1, Yokogawa) using solid-state lasers (395 nm, 488 nm, 561 nm) with appropriate filter sets (BFP: ET455/50M, GFP: ET525/36M, mCherry: ET605/52M) on 60× objectives (CFI PLAN APO OIL, NA=1.40, NIKON), using a CMOS camera (ORCA-Flash4.0). Photoactivation experiments used a digital micromirror device patterned illumination system (Polygon 4000, Photometrix). For simultaneous stimulation with 561 nm LED while imaging at 488ex/525em, we installed a custom dichroic TIRF filter cube (Chroma, ZT561DCRB-UF2) that permitted transmission of both shorter and longer wavelength light during 561nm LED stimulation. Cells were imaged in 35 mm imaging dishes (Matsunami, Dd35-14-1.5-U) within a stage-top environmental chamber (Tokai) at 37°C and 5% CO_2_. All microscope control and image analysis used Nikon NIS Elements AR software.

### Photoactivation Protocol

A small section of the cell was identified and illuminated using a user-defined region of interest (ROI) within Nikon NIS Elements AR Illumination Sequence Module. In all experiments, the Polygon400 and illumination/stimulation sequences were directly triggered by the camera. The standard illumination/stimulation sequence: 3 seconds of pHluorin (488 ex/525 em) pulsed acquisition (23 msec illumination every 50 msec) followed by 3 seconds of continuous 561 nm LED illumination to activate ArchT (power between 1% and 100%, see figure legends for details) with pHluorin (488 ex/525 em) pulsed co-acquisition (23 msec illumination every 50 msec). This pattern was repeated for a total of 154 seconds. For membrane dynamics experiments, an additional 30 sec of pHluorin (488 ex/525 em) pulsed acquisition was added both before and after the standard 154 sec pulsed protocol, for a total acquisition time of 215 sec. We note that we do observe some bleed-through of photomanipulation light into the pHluorin signal during the stimulation window. This produces an intensity “zigzag” pattern during photomanipulation with increased intensity measured in the 488ex/525em specifically during 561nm LED stimulation. This bleed-through is proportional to pHluorin intensity and occurs in both control and ArchT stimulated cells indicating true optical bleed-through and not a pH-dependent effect.

### LED titration and stimulation repeatability experiments

(Figure 2, S4) The LED power optimization and ArchT repeatability experiments were automated using the JOBs module within NIS Elements AR. For the power titration curves, the 154s illumination sequence (as described above) was looped with a rest period (no acquisition or illumination) of 2 minutes before the pattern repeated for the next LED power. For decreased power titration 100%, 50%, 20%, 10%, 1% LED power sequence was used. For increased power titration 10%, 20%, 30%, 50%, 100% sequence was used. For the repeatability assays, the illumination sequence pattern at 100% LED power was looped with a rest for 2 minutes. This illumination sequence pattern was performed 10 times on each cell.

**Figure 1.**
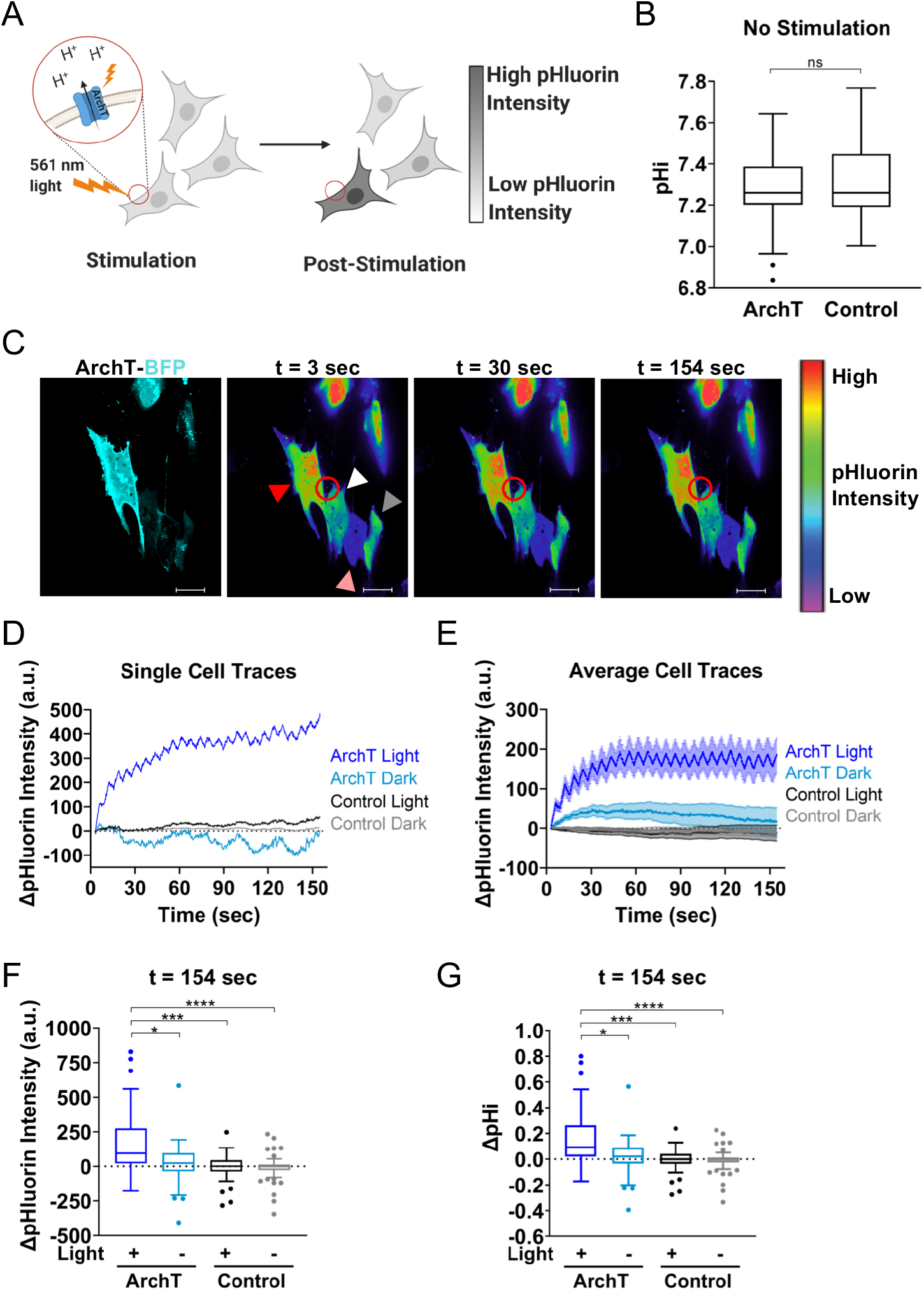
Archaerhodopsin can spatiotemporally increase pHi in single cells. **(A)** Spatially-restricted activation of ArchT-expressing cells by 561 nm light can raise pHi in a single cell and pH changes can be monitored using a genetically-encoded pH biosensor (pHluorin). **(B)** Quantification of resting pHi for RPE cells expressing ArchT compared to control RPE cells using a pH sensitive dye (see methods). Tukey boxplots (n = 51-98 cells per condition, 2 biological replicates). Significance determined using the Mann Whitney test. **(C)** An ArchT RPE cell (red arrow) and a control RPE cell (white arrow) are simultaneously photoactivated with 561 nm light within the stimulation region of interest (ROI, red circle). Also included in the field of view are unstimulated ArchT RPE cell (pink arrow) and an unstimulated control cell (grey arrow). Shown is an intensiometric display of pHluorin intensity during stimulation. Scale bar 20 μm. **(D)** Single-cell pHluorin intensity traces over time for cells in (C). **(E)** Quantification of pHluorin intensity changes for cells collected as described in (C). (n = 26-62, from 3-5 biological replicates), mean ± SEM. **(F)** Quantification data from (E) at the end of the experiment for stimulated (+ Light) and unstimulated (− Light) ArchT and control cells. Tukey boxplots. **(G)** Quantification of pHi changes in cells in (E) (see methods for details). Tukey boxplots. For F-G, significance determined using the Kruskal-Wallis test, Dunn’s multiple comparison correction, * p<0.05, ***p<0.001, **** p<0.0001.

**Figure 2.**
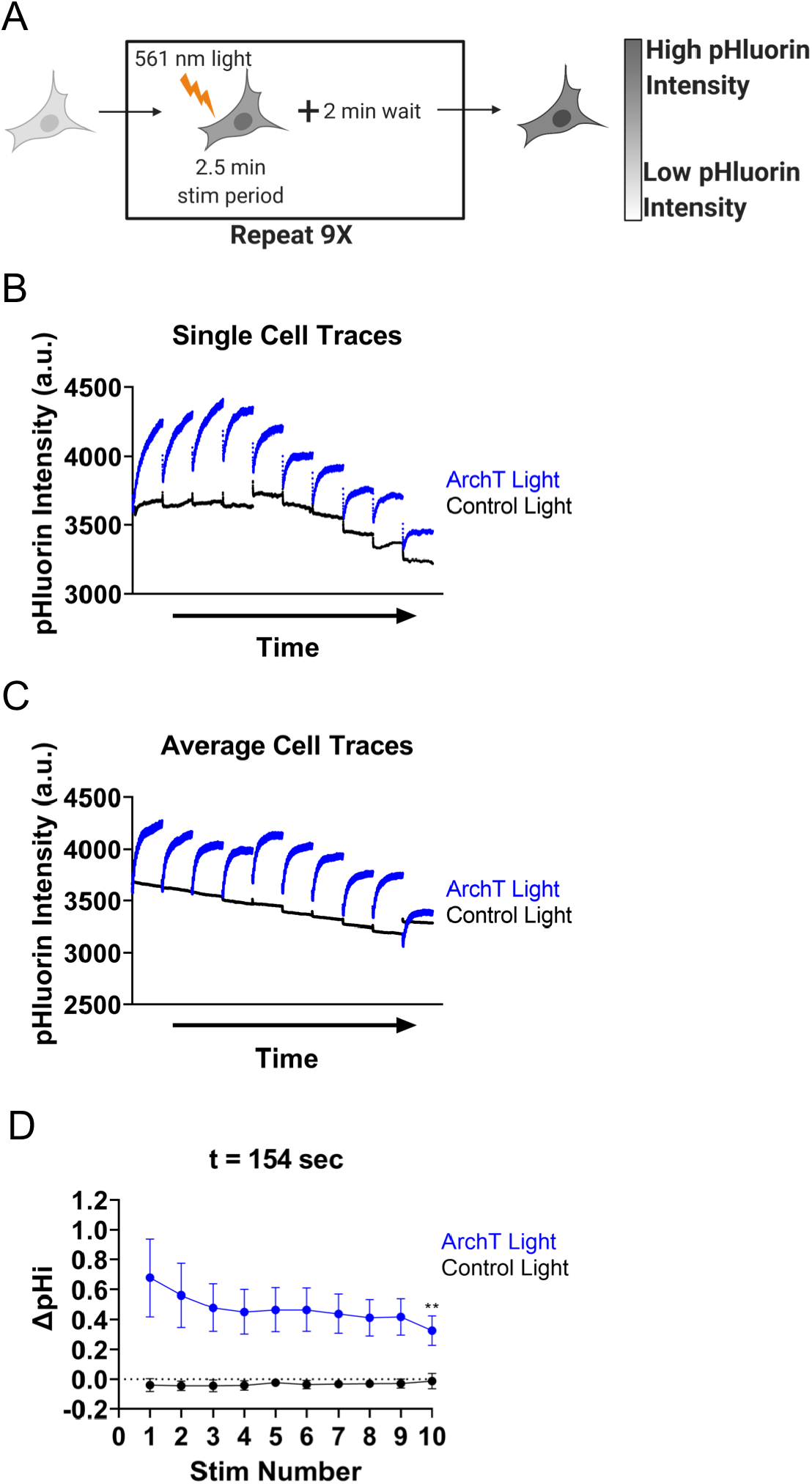
ArchT can be activated to repeatedly and robustly raise pHi in single cells. **(A)** Individual RPE cells expressing ArchT and pH biosensor (pHluorin) were stimulated using the standard protocol (see methods) followed by 2 minute recovery. This pattern is repeated 9 times. **(B)** Single-cell pHluorin intensity traces of single ArchT (blue) and Control (black) cells treated as described in A. **(C)** Average (mean) cell data collected as in B. (n=9-10 per condition, 3-4 biological replicates). **(D)** Change in pH intensity quantified for cells in (C), mean ± SEM. For (D), significance compared to Stim 1 determined using the two-way ANOVA, with Holm-Sidak multiple comparison correction, ** p<0.01.

### pHi Calculation using BCECF

(Figure 1B) The pHi of ArchT transfected RPE cells and control RPE cells were directly calculated using 2’,7’-bis(carboxyethyl)-5(6)-Carboxyfluorescein (BCECF, Biotium). BCECF is a ratiometric dual excitation/single-emission pH biosensor (405 ex/525 em for pH insensitive fluorescence, 488 ex/252 em for pH-sensitive fluorescence). Cells were loaded with 1 μM BCECF for 10 minutes, washed with complete media 3×5 minutes. We performed ratiometric imaging of cells loaded with BCECF dye (10 minutes, 37 C, 5% CO_2_). For standardization, pH standard buffers (pH ~6.5 and ~7.5 (0.025 M HEPES, 0.105 M KCl, 0.001 MgCl_2_)) were prepared with 10 μM nigericin (Invitrogen, N1495) (a protonophore) and added sequentially to cells to equilibrate pHe and pHi as previously described.^28^ Individual cell pHi was back-calculated using single-cell standard curves generated from ratiometric fluorescence in nigericin standard buffers and pHi values were reported in Figure 1B.

### pHi Calculation with ratiometric mCherry-pHluorin

(Figure S1) In Figure S1, pHi increases were directly calculated from pHluorin/mCherry ratios because as previously described^11^. Briefly, we took an initial acquisition of pHluorin/mCherry intensity ratios, then ran the stimulation protocol acquiring only in the pHluorin channel, then captured a post-stimulation acquisition of pHluorin/mCherry ratios. For standardization, pH standard buffers (pH ~6.5 and ~7.5 (0.025 M HEPES, 0.105 M KCl, 0.001 MgCl_2_)) were prepared with 10 μM nigericin (Invitrogen, N1495) and added to cells to equilibrate pHe and pHi as previously described.^28^ Individual cell pHi was back-calculated using single-cell standard curves and reported in Figure S1C.

### pHi Calculation from pHluorin intensity

(Figure 1, 2, S2, S3, S4, S5) For experiments where mCherry was not acquired, change in pHi was back-calculated from pHluorin intensity change using an average “standardization curve” from nigericin standardization datasets collected identically in both RPE and NIH-3T3 cells transfected with mCherry-pHluorin but not ArchT. Briefly, pHluorin/mCherry ratios were obtained for individual cells under standard culture conditions. Then pH standard buffers (pH ~6.5 and ~7.5 (0.025 M HEPES, 0.105 M KCl, 0.001 M MgCl_2_)) were prepared with 10 μM nigericin (Invitrogen, N1495) and added to cells to equilibrate pHe and pHi as previously described.^28^ For each cell, a standard curve was calculated using pHluorin/mCherry ratios at the high and low pHi standards. Using the average pHluorin intensity standard curve data from pHi measurements in RPE (see Figure S1A, outliers removed), and NIH-3T3 cells (see Figure S1B, outliers removed). We calculated the average intensity change in pHluorin per pH unit in each respective cell line, which was used to back-calculate change in pHi (Figure 1, 2, S2, S3, S4, S5) for each cell at 30 sec and 154 sec. Note that the NIH-3T3 standard curve was adjusted to account for differences in pHluorin exposure time between S1B (100 msec) and the stimulation experiments (50 msec).

### Data Analysis

Images were processed using Nikon Elements AR software. Images were background subtracted using an ROI placed on an area without visible cells within the field of view (Figures 1, 2, 3, S1, S2, S3, S4, and S5). ROIs were drawn on each cell using the pHluorin fluorescence signal and average ROI intensity values were obtained for pHluorin (pH-sensitive fluorescence) and mCherry (expression normalizer, where indicated in methods).

**Figure 3.**
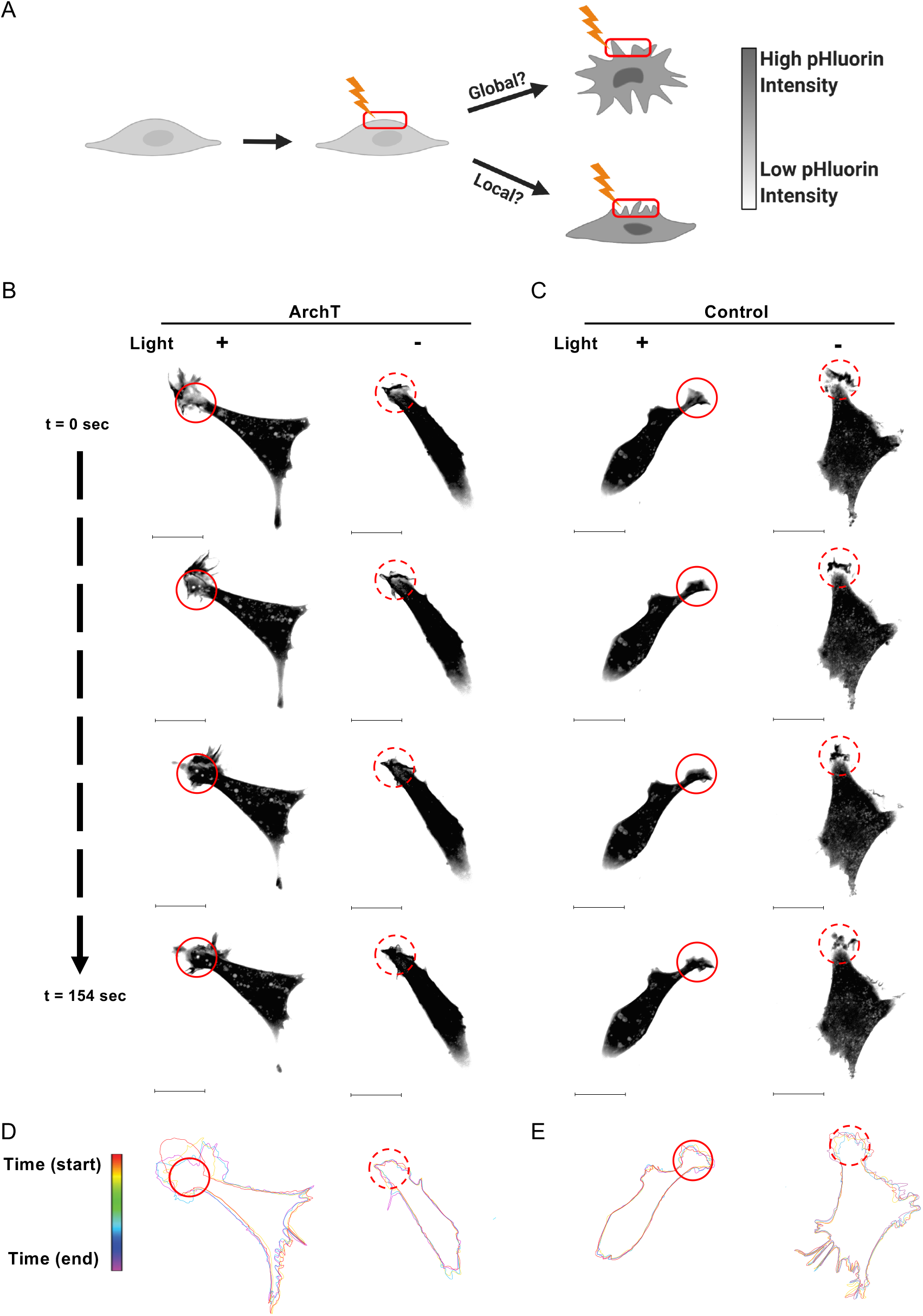
Increased pHi is a sufficient driver of local membrane protrusion events. **(A)** To better understand the role of pHi in driving membrane protrusion, the lamella of ArchT-expressing cells can be stimulated with 561 nm light and monitored for both global and local membrane ruffling events. **(B-C)** Representative images of ArchT (B) and Control (C) NIH-3T3 cells treated as depicted in A. The solid red circle indicates the stimulation ROI (+Light) while the dashed red circle indicates the mock-stim ROI (−Light). Stills from Movies S2-S5. Scale bar 20 μm. (**D-E**) For the ArchT (D) and control (E) cells depicted in B and C, membranes were traced at key video frames and overlaid: t = 0 sec (red) to 154 sec (violet). Additional representative cells can be found in Figure S6 (stimulated) and S7 (unstimulated).

### Photobleaching correction

(Figures 1, 2, S2, S3, S4, and S5). Briefly, average pHluorin ROI intensities of control cells were normalized to t = 0 and an average photobleaching curve was calculated from these individual curves. Data were corrected, with ArchT and control stimulated and unstimulated cells corrected with matched average stimulated and unstimulated photobleaching curves.

### Cell Traces

(Figures 3, S6, and S7). Adobe Illustrator was used to hand trace outlines of cells from key time frames within stimulation experiments (see representative videos provided: Supporting Movies 2-5).

### Localized membrane protrusion

(Figures 3, 4, 5, S6, S7, and S8). Note that cells were selected for stimulation by screening ArchT expression (for ArchT cells) and accessibility to a thin-lamella region to stimulate (for ArchT and control cells). All stimulated cells were processed through the analysis pipeline. A Nikon Elements AR General Analysis 3 (GA3) analysis pipeline was developed with the help of Nikon Instruments Inc. Software Support (Yu-Chen Hwang Ph.D., Biosystems Technical Solutions Manager). Briefly, each cell was segmented into 2 μm concentric segments originating from the circular stimulation ROI. For each concentric circle, total cell area and mean pHluorin intensity were determined within the segmented region. Change in area was tracked over time as a readout of local membrane protrusion events.

**Figure 4.**
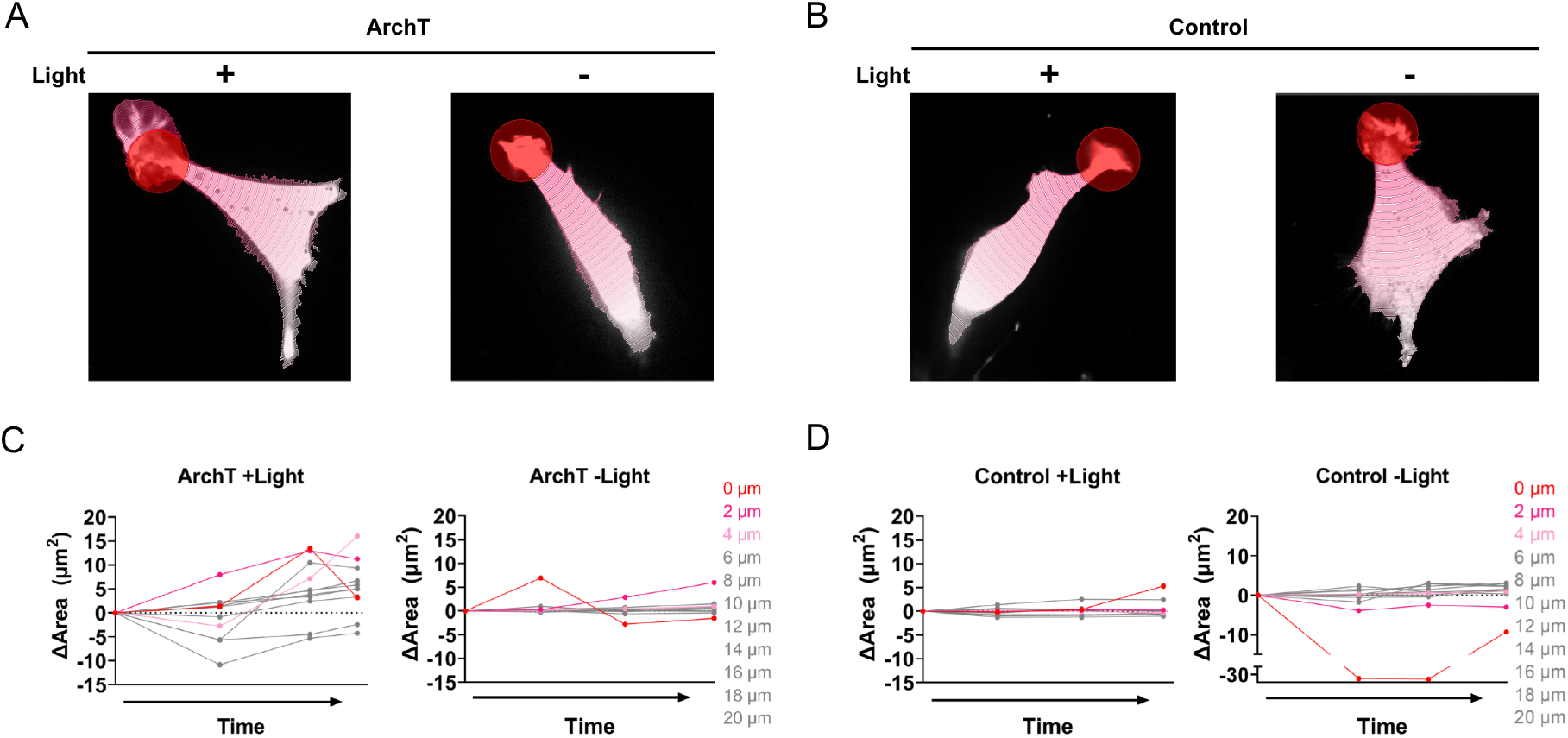
Quantification of local membrane protrusion events. **(A-B)** Representative segmentation analysis of ArchT (A) and Control (B) cells from Figure 3 (see methods). Briefly, concentric segmentation circles radiate out at 2 μm intervals from the edge of the stimulation (+Light) or mock stimulation (−Light) ROI (red circle). **(C-D)** Change in area for segmented ArchT (C) and Control (D) cells; where 0 μm indicates the stimulation ROI with increasing 2 μm intervals for each subsequent segment. Analysis of additional representative cells can be found in Figure S6 (stimulated) and S7 (unstimulated).

**Figure 5.**
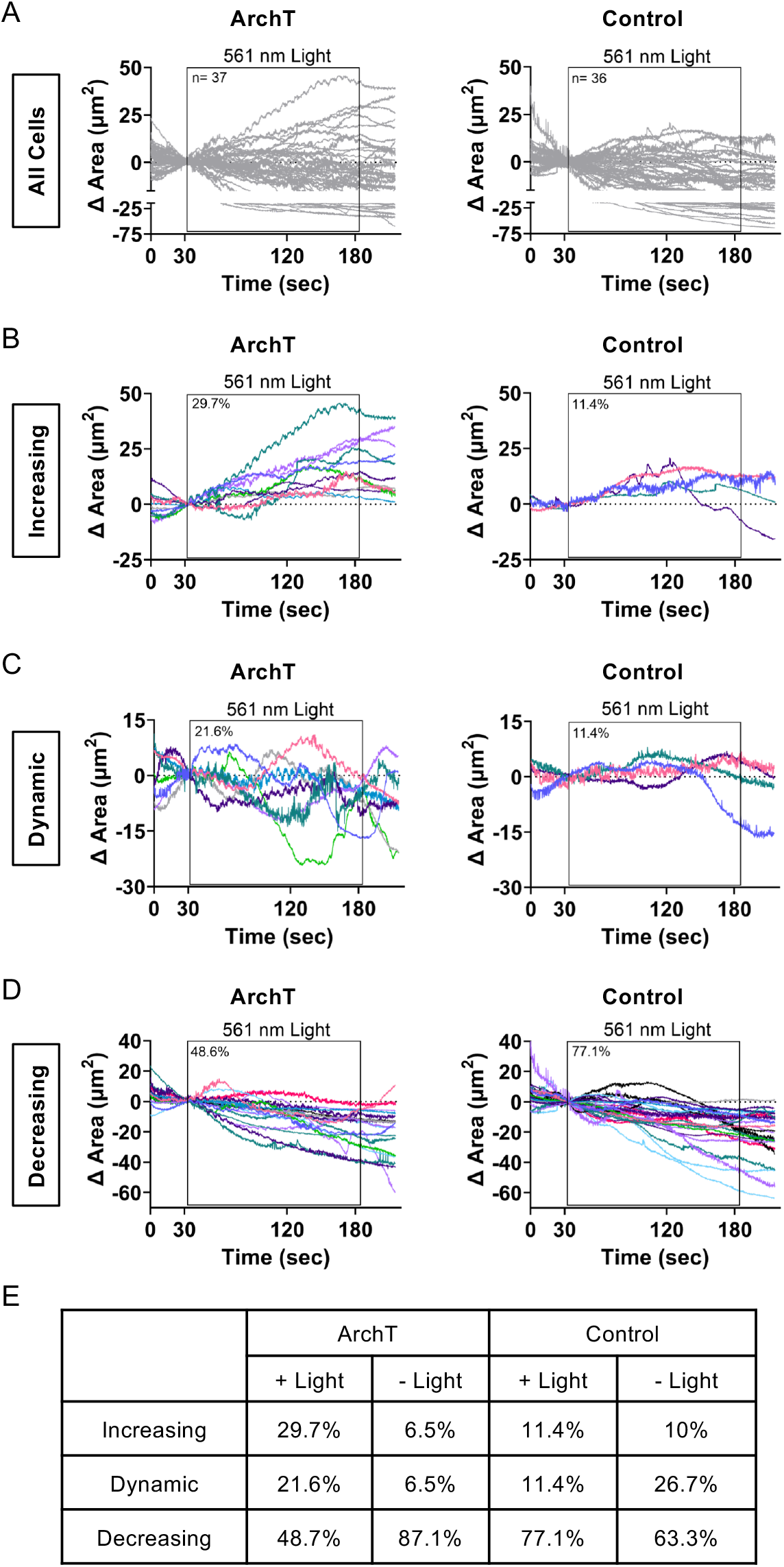
Increased pHi with ArchT induces increased membrane dynamics. **(A)** Membrane area changes were quantified within the 561 nm stimulation ROI for ArchT and control NIH-3T3 cells. Individual traces are shown for each cell, black box indicates timing of 561 nm light stimulation (n=36-37 cells per condition, from 5-6 biological replicates). **(B-D)** Binned traces of ArchT and control NIH-3T3 cells in **(A)** to characterize phenotypes that are protrusive **(B)**, dynamic **(C)**, and decreasing/static **(D)**. **(E)** Percentage of cells that fall under each phenotype (increasing, dynamic, or decreasing/static) for stimulated ArchT and Control cells (data shown in B-D) or mock-stimulated (−Light) ArchT and control cells (see Figure S8 for mock-stim data).

### Statistical Analysis

Normality of all of data was tested using Shapiro-Wilk normality test. For normally-distributed data (Figure 2), statistical significance was determined using a one-way ANOVA with Tukey multiple-comparisons correction. For non-normally distributed data (Figure 1, S1, S2, S3, and S5), statistical significance was determined using the Mann Whitney test for a single comparison or the Kruskal-Wallis test with Dunn’s multiple-comparisons correction for multiple comparisons. Significance for the LED power titration experiments (Figure S4) were determined using two-way ANOVA and Holm-Šídák’s multiple comparison test, and sphericity was assumed for these analyses.

## RESULTS

### ArchT induces spatiotemporal increases in pHi at the single-cell level

In our first validation experiment, we tested whether we could achieve spatiotemporal pHi increases in single human retinal epithelial (RPE) cells expressing ArchT (Figure 1A). We first confirmed good membrane localization of ArchT fused to blue fluorescent protein (BFP2) with an ER export signal (ArchT-BFP2-TSERex)^27^ (Figure S1A). We next tested whether ArchT expression alone altered resting pHi by measuring pHi in RPE cells transfected with ArchT compared to parental RPE cells using the pH sensitive dye 2’,7’-bis(carboxyethyl)-5(6)-Carboxyfluorescein (BCECF, see methods). Importantly, ArchT expression on its own does not increase resting pHi of cells, as ArchT transfected cells and control cells have the same back-calculated pHi (Figure 1B).

Unfortunately, BCECF photobleached too rapidly to allow for long-term pHi measurement. Thus, in order to monitor pHi changes in real time during spatiotemporal photomanipulation experiments, we co-transfected ArchT and a genetically encoded pH biosensor (mCherry-pHluorin)^29^. Using a digital micromirror device (DMD), we can spatially restrict 561 nm photoactivation to a single cell (Figure 1A). We developed a photomanipulation protocol that stimulates cells with 561 nm light in a spatially-restricted ROI using a 2.5 minute photomanipulation protocol (see methods). Briefly, we took an initial acquisition of pHluorin/mCherry fluorescence ratios, then ran the stimulation protocol to activate ArchT, then captured a post-stimulation acquisition of pHluorin/mCherry ratios. To back-calculate pHi from pHluorin/mCherry ratios as previously described^11,28^, we prepared pH standard buffers (pH ~6.5 and ~7.5) with 10 μM nigericin (a protonophore) and added to cells to equilibrate pHe and pHi.

If our tool and microscopy hardware enable spatiotemporal pHi manipulation, we would expect to observe increased pHi (increased pHluorin intensity) only in the ArchT-expressing stimulated cell (Figure 1A). Representative images in Figure S1B show simultaneous photoactivation in a selected region of interest of an ArchT-expressing cell and a control cell. We observed a robust increase in pHluorin/mCherry ratio only in the light-stimulated ArchT expressing cell, while there was no change in pHluorin/mCherry ratio in the light-stimulated control cell (mCherry-pHluorin only) (Figure S1B, Movie S1). When this analysis is performed across many cells, we found that ArchT stimulated cells increased pHi by an average of 0.31 ± 0.06 (mean ± SEM) pH units by the end of the experiment while the pHi of control cells was unaffected at the end of the stimulation protocol (Figure S1C). However, ratiometric pHluorin/mCherry pHi quantification requires 561 illumination across the whole field of view, which could serve to pre-activate ArchT cells outside of the designated stimulation ROI. Indeed, we did observe significant pHi increases in unstimulated ArchT cells in the same fields of view (0.17 ± 0.03 pH units, mean ± SEM) (Figure S1C). This result suggests that even short illumination with low power 561 nm laser light can activate ArchT, producing a high level of apparent ArchT “dark activation” in unstimulated ArchT cells when using this experimental protocol.

In order to reduce dark activation, we optimized our stimulation protocol to eliminate mCherry acquisition to avoid activating ArchT prior to stimulation. Instead, we monitored only pHluorin intensity during the 2.5 minute photomanipulation protocol and applied an average nigericin standard curve to convert change in pHluorin signal to change in pHi (Figure S2A, see methods for details). Representative images in Figure 1C show simultaneous photoactivation in a selected region of interest (red circle) of an ArchT-expressing cell (red arrow) and a control cell (white arrow). We observed a robust increase in pHluorin intensity in the light-stimulated cell, while there was no net change in pHluorin intensity in control simulated cells (white arrow) (Figure 1C and D). We also observed no net change in pHluorin intensity in unstimulated ArchT cells (pink arrow) or unstimulated control cells (grey arrow) within the same field of view (Figure 1C and D). When this analysis is performed across many ArchT cells, we observed a rapid increase in pHluorin intensity within just 30 seconds that plateaus by the end of the assay (Figure 1E). Unstimulated ArchT cells did not exhibit a significant increase in pHluorin fluorescence, indicating spatially restricted photoactivation (Figure 1E and F, Figure S2B (30 sec)). Importantly, pHluorin signal did not increase in control cells, regardless of whether they were stimulated with light (Figure 1E and F, Figure S2B (30 sec)). Thus, the pHluorin response requires both expression of the ArchT tool and light activation.

We quantified pHi increases induced by ArchT in single cells using only the pHluorin intensity values (see methods). We found that pHi increased in ArchT stimulated cells by an average of 0.15 ± 0.03 (mean ± SEM) pH units after 30 seconds (Figure S2C) and 0.18 ± 0.04 (mean ± SEM) pH units by the end of the experiment (Figure 1G). At the end of the stimulation period, stimulated ArchT cells were the only cells with statistically increased pHi (compared to zero, p<0.0001). This result suggests that dark activation of ArchT is minimized using this updated stimulation protocol that avoids any pre-exposure to 561 nm light. Importantly, the specific pHi increases achieved only in ArchT stimulated cells correspond well with physiological pHi increases (0.1-0.35) observed during normal cell behaviors like directed cell migration^1,2,5,28^, cell polarization^1,2^, and cell cycle progression^30^.

### ArchT induces robust pHi increases at low and high stimulation powers

We next sought to determine the range of LED stimulation powers that can be used to induce a robust pHi increase. We note that when using the strongest LED power (100%) in RPEs we would expect both heat effects and potential for ArchT dark activation to be high. When using 100% LED power in RPEs (Figure S3) we saw a similar pHi increase for ArchT cells (0.10 ± 0.04), at the end of the experiment, compared to the lower 30% laser power experiments (Figure 1). At 100% stimulation power, we did observe increased ArchT dark-activation, leading to both stimulated and unstimulated ArchT cells demonstrating an increase in pHi (compared to zero, p<0.05) at the end of the experiment. This likely reflects increased light bleed-through outside the stimulation ROI at 100% LED power, leading to increased measured “dark activation” of ArchT under these conditions. We saw no increase in pHi in stimulated control cells, indicating heat effects alone are not sufficient to induce pHi changes observed in these assays. These data suggest that even the highest LED stimulation settings produce robust and specific increases in pHi in ArchT stimulated cells compared to control stimulated cells (Figure S3).

If ArchT could be specifically activated to raise pHi using a range of LED stimulation powers, the tool would be adaptable to various experimental applications. To test this, we first titrated LED power sequentially up from 10% to 100% LED power and observed statistically significant increases in pHi compared to control at as little as 10% power (Figure S4A, B). We also titrated LED power sequentially down from high intensity (100%) to low intensity (1%) to control for potential prolonged heat effects producing a different outcome on control cell responses. In this case, we found that pHi was increased compared to control as low as 20% LED power (Figure S4C, D). At the end of stimulation period, we observe no significant differences in pHi increases achieved with LED stimulation ranging from 1% to 100%. We also observe significant increases in pHi (compared to zero, p<0.0001) within ArchT stimulated cells for stimulation powers as low as 10% whether titrating up or down in power. This shows that the ArchT tool can be stimulated using a range of LED powers to fit the imaging needs of the user. In particular, these results show that lower LED stimulation powers can be used to significantly raise pHi if phototoxicity, photobleaching, dark activation, or heat-effects are a concern in sensitive applications. Collectively, these data demonstrate that ArchT can be used to elicit robust pHi responses under various LED stimulation conditions.

### ArchT-induced pHi increases are repeatable within single cells

We have shown that the ArchT tool can be used to increase pHi in single-cells in real time and we observe efficient and robust increases in pHi using a range of LED stimulation powers. Next, we determined whether the ArchT tool can be used to repeatedly and reversibly increase pHi in a single cell. Achieving repeatable pHi manipulations will enable investigation of pH-dependent cell behaviors that occur on longer timescales, such as cell polarization and migration. As a proof of principle, we stimulated an RPE cell expressing ArchT and the pH biosensor with 561 nm light using the optimized 2.5-minute stimulation protocol (see methods), and then we allowed the cell to recover for 2 minutes. This pattern was repeated for a total of 10 stimulations on the same individual targeted cell (Figure 2A). As expected, within each stimulation period we observed a rapid increase in pHluorin intensity in the light-stimulated ArchT cell, followed by a plateauing of the response, while the control cells were non-responsive (Figure 2B). However, between stimulation periods, pHluorin intensity recovered to baseline levels, suggesting that the cells were returning to pHi homeostasis during the rest period (Figure 2B). This result was robust across many ArchT and control cells over the 45-minute repeated stimulation protocol (Figure 2C). Notably, the measured increases in pHi in ArchT cells during stimulation windows were consistent and robust (Figure 2D). These data suggest that the ArchT tool can be used to repeatedly and reliably raise pHi in cells over a 45-minute protocol. Furthermore, these results suggest that ArchT may be an appropriate tool to investigate pH-dependent cell behaviors, such as cell polarization, that occur on a longer timescale.^31^

### Increases in pHi are sufficient to drive localized membrane protrusion

We next applied ArchT to investigate the relationship between spatiotemporally increased pHi and single-cell membrane ruffling responses, which occur on the order of seconds to minutes^3,4,32^. The ArchT tool allows us to determine whether localized pHi increases produce global or localized membrane ruffling responses in single cells (Figure 3A). One hypothesis is that protons are diffusible and thus a local pHi increase would quickly produce a global (non-localized) membrane ruffling response.^33^ An alternative hypothesis is that localized increases in pHi could be maintained or reinforced by protein recruitment^34^ to produce localized membrane ruffling responses. With ArchT as a tool for spatiotemporal pHi manipulation, we can now determine if increased pHi is sufficient for membrane remodeling in single-cells. We selected mouse fibroblasts NIH-3T3 (NIH-3T3) cells because they form more pronounced lamella than RPE cells, making them ideal for investigating the relationship between increased pHi and membrane ruffling. We first validated that ArchT could produce similar pHi increases in NIH-3T3 cells (Figure S5) and observed comparable specificity and pHi changes to those we observed in RPE cells. At the end of the stimulation period, we observe significant increases in ArchT stimulated cells (compared to 0, p<0.01), but not in ArchT unstimulated cells or control cells. This suggests that ArchT can be used to produce robust physiological and spatiotemporal increases in pHi in single NIH-3T3 cells.

We expressed ArchT in NIH-3T3 cells and stimulated the thin-lamella portions of the cell with 561 nm light. We selected fields of view based on expression of ArchT and the presence of an accessible thin lamella. For control cells (mCherry-pHluorin expression alone) we similarly selected cells based on the ability to target a thin lamella. In order to confirm that ArchT cells are not inherently more dynamic than control cells, we included mock-stimulation experiments where membrane dynamics were monitored over 2.5 minutes but no stimulation light was applied to the cells. When the thin-lamella region of ArchT cells was stimulated with light, we observed distinct and localized membrane ruffling around the stimulation ROI (Figure 3B, Movie S2). This effect was not observed in control stimulated cells or in unstimulated ArchT or control cells (Figure 3B and C, Movie S3 (ArchT –Light), Movie S4 (Control +Light), and Movie S5 (Control –Light)).

In order to better analyze this response, we traced the outline of the cells at various time points throughout the experiment. From these traces, we can see that the stimulated ArchT cell is more dynamic in the stimulation region compared to the unstimulated ArchT cell (Figure 3D) and both stimulated and unstimulated control cells (Figure 3E). The ArchT cell starts with a pronounced lamella protrusion at the top-left of the cell (Figure 3D, red trace) and ends with the protrusion extended above and to the bottom right of the stimulation ROI (Figure 3D, purple trace). Additional examples of ArchT and control cell traces can be found in Figure S6 (stimulated) and S7 (unstimulated). A lack of membrane dynamics outside the stimulated ROI in the ArchT cell suggests a localized membrane response in or near the stimulation ROI. This localized stimulation region response is unlikely to be driven by heat or light as stimulated control cells lack any similar membrane dynamics in the stimulation region (Figure 3E, Figure S6). These data support the hypothesis that spatiotemporal increases in pHi are sufficient to drive localized membrane ruffling responses in single cells.

To quantify localized pHi-dependent membrane ruffling responses, we developed a custom analysis within NIS Elements (see methods). Briefly, each cell was segmented into a series of 2 μm wide concentric circles, originating from the circular stimulation region (Figure 4A-B). This segmentation allowed us to monitor membrane ruffling (change in cell area) as a function of distance from stimulation region. We observed that for ArchT stimulated cells, there was a distinct distance correlation (Figure 4C, additional representative cells in Figure S6), with the stimulation region (0 μm, red trace) and immediate neighbor regions (2–4 μm, pink traces) having a larger net change in area compared to more distant regions (6–20 μm, grey traces). Notably, unstimulated ArchT cells (Figure 4A,C) as well as stimulated and unstimulated control cells (Figure 4B,D) are less dynamic (smaller area change) and there is no observed distance-dependence to the stimulation ROI in any of the controls (Additional examples in Figure S6 (stimulated cells) and Figure S7 (unstimulated cells)). Thus these membrane ruffling events require both expression and light-activation of ArchT.

In our initial analyses of these pH-induced membrane dynamics, we noted that some cells had pronounced localized protrusions with spatiotemporal increases in pHi, while other cells exhibited both protrusion and retraction events. As these events would be lost in our previous analyses of net area change, we sought to quantify the *dynamics* of membrane responses in ArchT versus control. We modified the standard stimulation protocol to include a 30 sec observation period before and after the stimulation period (see methods), allowing us to better compare membrane ruffling during stimulation to basal dynamics within the exact same cell.

To investigate these dynamics more closely, we quantified area change within the stimulation ROI for each stimulated (Figure 5A) and mock-stimulated cell (Figure S8A). From this quantification, we identified 3 distinct response phenotypes: increasing protrusive responses (over 10 μm^2^ area increase compared to starting point), dynamic responses (dynamic change in area of at least 10 μm^2^ and crossed the x-axis at least 3 times), and decreasing retraction responses (over 10 μm^2^ area decrease compared to starting point). We binned all cells by response, and found that 29.7% of ArchT stimulated cells exhibited a strong protrusive response (increase in area) during the photo-stimulation period while only 11.4% of stimulated control cells fell into this protrusive category (Figure 5B, E). Only 6.5% of unstimulated ArchT cells fell into this category (Figure 5E and S8B) further supporting the role of ArchT activation-dependent pHi increases in driving a protrusive phenotype. We note that the protrusive phenotype was smaller and light-independent in control stimulated (Figure 5B) and unstimulated ArchT and control cells (Figure S8B). These results indicate that our stimulation protocol is not inhibiting naturally occurring membrane protrusion events as control stimulated and unstimulated cells have a similar rates of protrusion (Figure 5E). Furthermore, increased protrusion in ArchT stimulated cells support the conclusion that increased pHi is a sufficient driver of localized and sustained membrane protrusion.

A large proportion of stimulated ArchT cells (21.6%) exhibited dynamic area changes during the photo-stimulation period compared to just 6.5% of unstimulated ArchT cells (Figure 5C, E and Figure S8C). Again, while some stimulated and unstimulated control cells exhibited dynamic responses, the magnitude of responses was attenuated compared to stimulated ArchT cells (Figure 5C, E and Figure S8C). We note that a larger percentage of mock-stimulated control cells exhibited dynamic protrusions compared to the stimulated control Cells (Figure E). This may suggest that stimulation light reduces dynamic ruffling, but it could also be an artifact of arbitrary definition of “stim” region for the mock-stim conditions. Importantly, the distinct light-dependent increases in both protrusive and dynamic membrane ruffling phenotypes in ArchT cells are not observed in control cells. This suggests a role for pHi increases in driving membrane dynamics as well as sustained protrusion. For cells with retraction responses during the stimulation period, the ArchT and control cells had similar responses (Figure 5D, E). Unlike the other ArchT responses described, we note that retraction responses appear to be independent of stimulation light, with measured retraction prior to the stimulation window as well as during stimulation (Figure 5D and Figure S8D). This may suggest that depolymerization of actin fibers during retraction events dominate pHi-dependent protrusion events. Taken together, these data suggest that ArchT induces localized pHi increases which in turn induce localized and dynamic membrane ruffling. Furthermore, our data suggest that photoactivation-dependent ruffling dynamics in ArchT cells are induced specifically by increased pHi.

## DISCUSSION

Current approaches to manipulate pHi lack spatiotemporal control, limiting our understanding of the role of pHi dynamics in driving cellular processes. Furthermore, reliance on population-level analyses can obscure a role for pHi in behavioral or phenotypic cellular heterogeneity. In this work, we have shown that the light-activated proton pump ArchT can be used as a robust optogenetic tool to spatiotemporally increase pHi in single-cells. The tool can be used to increase pHi over short time periods (minutes) and can be repeatedly stimulated to increase pHi for a longer period of time (~45 minutes). Using this tool, we show that spatially-restricted activation of ArchT increases pHi and drives localized pHi-dependent membrane ruffling. Our current developed protocols will allow us to apply ArchT to investigate roles for pHi in regulating more complex single-cell behaviors such as cell polarization and migration.

Future work will further investigate the dynamic membrane ruffling observed within ArchT cells with the goal of determining the molecular determinants of these responses. One caveat to these results is that Archaerhodopsins have been shown to hyperpolarize the cell membrane,^24,25,35^ and previous work has linked membrane potential changes, both depolarization and hyperpolarization, to cytoskeleton remodeling on the timescale of 5-30 minutes^36,37^. Future work will be required to fully decouple effects of membrane polarization and pHi dynamics on the phenotypes reported here. One key aspect of this future work will be the development of a light-activatable electroneutral exchanger, such as the sodium-proton exchanger (NHE1), that would allow pHi increases to be decoupled from membrane potential changes.

The work described herein provides an experimental platform to transform our understanding of how pHi dynamics regulate normal cell behaviors. However, dysregulated pHi dynamics are a hallmark of diseases such as cancer (constitutively increased pHi)^6–8^ and is thought to be an early event in cancer development.^11,38^ Using ArchT to spatiotemporally increase pHi in single cells will allow us to probe whether increased pHi is a sufficient driver of single cell behaviors and whether increasing pHi in a single cell in an epithelial layer results in pHi communication to neighboring cells. With our development of these ArchT pHi manipulation protocols, these complex but critical questions of basic and cancer cell biology are now within our experimental grasp.

## Acknowledgments

We would like to thank Dr. Yu-chen Hwang and Nikon Instruments Inc. Software Support Group for assistance developing the custom NIS Elements analysis pipeline. We would also like to thank members of the White lab for their helpful feedback on the manuscript. Figure 1A, 2A, and 3A created with BioRender.com.

## List of Supporting Information

Supporting Information includes Figures S1-S8 and Movies S1-S5. Figures include additional characterization of ArchT in RPE cells and in NIH-3T3 cells, ArchT responses to various LED stimulation power, additional cell traces as in Figure 3, and additional characterization of mock-stimulation cell membrane dynamics (similar to that described in Figure 5). Movies show ArchT stimulation and pHi response as well as ArchT and control membrane dynamics response.

Supporting Figures 1-8

Supporting Movies 1-5

Movie S1: ArchT RPE (red arrow) and control RPE cells (white arrow) expressing mCherry-pHluorin are stimulated with 561 nm light in a region of interest (red circle). The pHluorin intensity increases in the ArchT stimulated cell, but not in the control stimulated cell. Stills from video shown in Figure S1.

Movie S2: Lamella of ArchT RPE cell expressing mCherry-pHluorin is stimulated with 561 nm light within a region of interest (red circle, see methods for details). A localized membrane ruffling response is observed within and directly adjacent to the stimulation region. Stills from video shown in Figure 3 and quantified in Figure 4.

Movie S3: Lamella of ArchT RPE cell expressing mCherry-pHluorin is mock-stimulated within a region of interest (red circle, see methods for details). No localized membrane responses are observed. Stills from video shown in Figure 3 and quantified in Figure 4.

Movie S4: Lamella of control RPE cell expressing mCherry-pHluorin is stimulated with 561 nm light within a region of interest (red circle, see methods for details). No localized membrane responses are observed. Stills from video shown in Figure 3 and quantified in Figure 4.

Movie S5. Lamella of control RPE cell expressing mCherry-pHluorin is mock-stimulated within a region of interest (red circle, see methods for details). No localized membrane responses are observed. Stills from video shown in Figure 3 and quantified in Figure 4.

## For Table of Contents Only

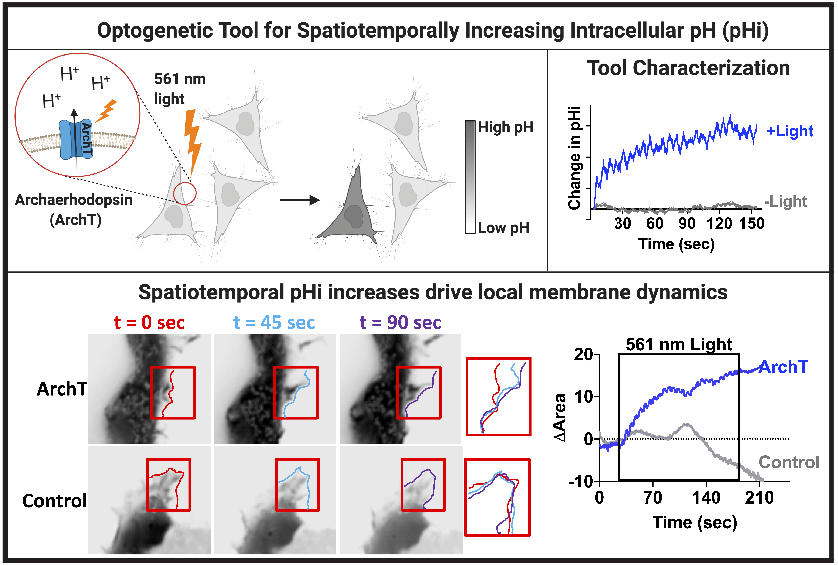

## References

(1) Frantz, C.; Karydis, A.; Nalbant, P.; Hahn, K. M.; Barber, D. L. Positive Feedback between Cdc42 Activity and H ^+^ Efflux by the Na-H Exchanger NHE1 for Polarity of Migrating Cells. The Journal of Cell Biology 2007, 179 (3), 403–410. https://doi.org/10.1083/jcb.200704169.

(2) Denker, S. P.; Barber, D. L. Cell Migration Requires Both Ion Translocation and Cytoskeletal Anchoring by the Na-H Exchanger NHE1. The Journal of Cell Biology 2002, 159 (6), 1087–1096. https://doi.org/10.1083/jcb.200208050.

(3) Frantz, C.; Barreiro, G.; Dominguez, L.; Chen, X.; Eddy, R.; Condeelis, J.; Kelly, M. J. S.; Jacobson, M. P.; Barber, D. L. Cofilin Is a PH Sensor for Actin Free Barbed End Formation: Role of Phosphoinositide Binding. J. Cell Biol. 2008, 183 (5), 865–879. https://doi.org/10.1083/jcb.200804161.

(4) Srivastava, J.; Barreiro, G.; Groscurth, S.; Gingras, A. R.; Goult, B. T.; Critchley, D. R.; Kelly, M. J. S.; Jacobson, M. P.; Barber, D. L. Structural Model and Functional Significance of PH-Dependent Talin-Actin Binding for Focal Adhesion Remodeling. Proceedings of the National Academy of Sciences 2008, 105 (38), 14436–14441. https://doi.org/10.1073/pnas.0805163105.

(5) Martin, C.; Pedersen, S. F.; Schwab, A.; Stock, C. Intracellular PH Gradients in Migrating Cells. American Journal of Physiology-Cell Physiology 2010, 300 (3), C490–C495. https://doi.org/10.1152/ajpcell.00280.2010.

(6) Webb, B. A.; Chimenti, M.; Jacobson, M. P.; Barber, D. L. Dysregulated PH: A Perfect Storm for Cancer Progression. Nat. Rev. Cancer 2011, 11 (9), 671–677. https://doi.org/10.1038/nrc3110.

(7) Czowski, B. J.; Romero-Moreno, R.; Trull, K. J.; White, K. A. Cancer and PH Dynamics: Transcriptional Regulation, Proteostasis, and the Need for New Molecular Tools. Cancers (Basel) 2020, 12 (10). https://doi.org/10.3390/cancers12102760.

(8) White, K. A.; Grillo-Hill, B. K.; Barber, D. L. Cancer Cell Behaviors Mediated by Dysregulated PH Dynamics at a Glance. J. Cell. Sci. 2017, 130 (4), 663–669. https://doi.org/10.1242/jcs.195297.

(9) Majdi, A.; Mahmoudi, J.; Sadigh-Eteghad, S.; Golzari, S. E. J.; Sabermarouf, B.; Reyhani-Rad, S. Permissive Role of Cytosolic PH Acidification in Neurodegeneration: A Closer Look at Its Causes and Consequences. Journal of Neuroscience Research 2016, 94 (10), 879–887. https://doi.org/10.1002/jnr.23757.

(10) Harguindey, S.; Stanciu, D.; Devesa, J.; Alfarouk, K.; Cardone, R. A.; Polo Orozco, J. D.; Devesa, P.; Rauch, C.; Orive, G.; Anitua, E.; Roger, S.; Reshkin, S. J. Cellular Acidification as a New Approach to Cancer Treatment and to the Understanding and Therapeutics of Neurodegenerative Diseases. Semin Cancer Biol 2017, 43, 157–179. https://doi.org/10.1016/j.semcancer.2017.02.003.

(11) Grillo-Hill, B. K.; Choi, C.; Jimenez-Vidal, M.; Barber, D. L. Increased H+ Efflux Is Sufficient to Induce Dysplasia and Necessary for Viability with Oncogene Expression. eLife 2015, 4, e03270. https://doi.org/10.7554/eLife.03270.

(12) Parks, S. K.; Cormerais, Y.; Durivault, J.; Pouyssegur, J. Genetic Disruption of the PHi-Regulating Proteins Na+/H+ Exchanger 1 (SLC9A1) and Carbonic Anhydrase 9 Severely Reduces Growth of Colon Cancer Cells. Oncotarget 2016, 8 (6), 10225–10237. https://doi.org/10.18632/oncotarget.14379.

(13) Slepkov, E. R.; Rainey, J. K.; Sykes, B. D.; Fliegel, L. Structural and Functional Analysis of the Na+/H+ Exchanger. Biochem J 2007, 401 (Pt 3), 623–633. https://doi.org/10.1042/BJ20061062.

(14) Stock, C.; Schwab, A. Role of the Na/H Exchanger NHE1 in Cell Migration. Acta Physiol (Oxf) 2006, 187 (1–2), 149–157. https://doi.org/10.1111/j.1748-1716.2006.01543.x.

(15) Olesen, C. W.; Vogensen, J.; Axholm, I.; Severin, M.; Schnipper, J.; Pedersen, I. S.; von Stemann, J. H.; Schrøder, J. M.; Christensen, D. P.; Pedersen, S. F. Trafficking, Localization and Degradation of the Na + ,HCO 3 − Co-Transporter NBCn1 in Kidney and Breast Epithelial Cells. Scientific Reports 2018, 8 (1), 7435. https://doi.org/10.1038/s41598-018-25059-7.

(16) Rolver, M. G.; Elingaard-Larsen, L. O.; Andersen, A. P.; Counillon, L.; Pedersen, S. F. Pyrazine Ring-Based Na+/H+ Exchanger (NHE) Inhibitors Potently Inhibit Cancer Cell Growth in 3D Culture, Independent of NHE1. Sci Rep 2020, 10. https://doi.org/10.1038/s41598-020-62430-z.

(17) Mauvezin, C.; Nagy, P.; Juhász, G.; Neufeld, T. P. Autophagosome–Lysosome Fusion Is Independent of V-ATPase-Mediated Acidification. Nature Communications 2015, 6 (1), 7007. https://doi.org/10.1038/ncomms8007.

(18) Yuan, Y.; Shimura, M.; Hughes, B. A. Regulation of Inwardly Rectifying K+ Channels in Retinal Pigment Epithelial Cells by Intracellular PH. The Journal of Physiology 2003, 549 (2), 429–438. https://doi.org/10.1113/jphysiol.2003.042341.

(19) Grinstein, S.; Romanek, R.; Rotstein, O. D. Method for Manipulation of Cytosolic PH in Cells Clamped in the Whole Cell or Perforated-Patch Configurations. Am J Physiol 1994, 267 (4 Pt 1), C1152–1159. https://doi.org/10.1152/ajpcell.1994.267.4.C1152.

(20) Ellis-Davies, G. C. R. Caged Compounds: Photorelease Technology for Control of Cellular Chemistry and Physiology. Nat Methods 2007, 4 (8), 619–628. https://doi.org/10.1038/nmeth1072.

(21) Kim, J. H.; Demaurex, N.; Grinstein, S. Chapter 20 Intracellular PH: Measurement, Manipulation and Physiological Regulation. In Handbook of Biological Physics; Konings, W. N., Kaback, H. R., Lolkema, J. S., Eds.; Transport Processes in Eukaryotic and Prokaryotic Organisms; North-Holland, 1996; Vol. 2, pp 447–472. https://doi.org/10.1016/S1383-8121(96)80061-3.

(22) Hulikova, A.; Swietach, P. Nuclear Proton Dynamics and Interactions with Calcium Signaling. Journal of Molecular and Cellular Cardiology 2016, 96, 26–37. https://doi.org/10.1016/j.yjmcc.2015.07.003.

(23) Miesenböck, G. Optogenetic Control of Cells and Circuits. Annu Rev Cell Dev Biol 2011, 27, 731–758. https://doi.org/10.1146/annurev-cellbio-100109-104051.

(24) Chow, B. Y.; Han, X.; Dobry, A. S.; Qian, X.; Chuong, A. S.; Li, M.; Henninger, M. A.; Belfort, G. M.; Lin, Y.; Monahan, P. E.; Boyden, E. S. High-Performance Genetically Targetable Optical Neural Silencing by Light-Driven Proton Pumps. Nature 2010, 463 (7277), 98–102. https://doi.org/10.1038/nature08652.

(25) Han, X.; Chow, B. Y.; Zhou, H.; Klapoetke, N. C.; Chuong, A.; Rajimehr, R.; Yang, A.; Baratta, M. V.; Winkle, J.; Desimone, R.; Boyden, E. S. A High-Light Sensitivity Optical Neural Silencer: Development and Application to Optogenetic Control of Non-Human Primate Cortex. Front. Syst. Neurosci. 2011, 5. https://doi.org/10.3389/fnsys.2011.00018.

(26) El-Gaby, M.; Zhang, Y.; Wolf, K.; Schwiening, C. J.; Paulsen, O.; Shipton, O. A. Archaerhodopsin Selectively and Reversibly Silences Synaptic Transmission through Altered PH. Cell Rep 2016, 16 (8), 2259–2268. https://doi.org/10.1016/j.celrep.2016.07.057.

(27) Wu, L.; Dong, A.; Dong, L.; Wang, S.-Q.; Li, Y. PARIS, an Optogenetic Method for Functionally Mapping Gap Junctions. eLife 8. https://doi.org/10.7554/eLife.43366.

(28) Choi, C.-H.; Webb, B. A.; Chimenti, M. S.; Jacobson, M. P.; Barber, D. L. PH Sensing by FAK-His58 Regulates Focal Adhesion Remodeling. The Journal of Cell Biology 2013, 202 (6), 849–859. https://doi.org/10.1083/jcb.201302131.

(29) Koivusalo, M.; Welch, C.; Hayashi, H.; Scott, C. C.; Kim, M.; Alexander, T.; Touret, N.; Hahn, K. M.; Grinstein, S. Amiloride Inhibits Macropinocytosis by Lowering Submembranous PH and Preventing Rac1 and Cdc42 Signaling. Journal of Cell Biology 2010, 188 (4), 547–563. https://doi.org/10.1083/jcb.200908086.

(30) Flinck, M.; Kramer, S. H.; Pedersen, S. F. Roles of PH in Control of Cell Proliferation. Acta Physiologica 2018, 223 (3), e13068. https://doi.org/10.1111/apha.13068.

(31) Rappel, W.-J.; Edelstein-Keshet, L. Mechanisms of Cell Polarization. Curr Opin Syst Biol 2017, 3, 43–53. https://doi.org/10.1016/j.coisb.2017.03.005.

(32) Gov, N. S.; Gopinathan, A. Dynamics of Membranes Driven by Actin Polymerization. Biophys J 2006, 90 (2), 454–469. https://doi.org/10.1529/biophysj.105.062224.

(33) Propagation of Protons at the Water Membrane Interface Microscopic Evaluation of a Macroscopic Process; CRC Press, 2017; pp 259–276. https://doi.org/10.1201/9780203743805-12.

(34) Wu, Y. I.; Frey, D.; Lungu, O. I.; Jaehrig, A.; Schlichting, I.; Kuhlman, B.; Hahn, K. M. A Genetically-Encoded Photoactivatable Rac Controls the Motility of Living Cells. Nature 2009, 461 (7260), 104–108. https://doi.org/10.1038/nature08241.

(35) Adams, D. S.; Tseng, A.-S.; Levin, M. Light-Activation of the Archaerhodopsin H+-Pump Reverses Age-Dependent Loss of Vertebrate Regeneration: Sparking System-Level Controls in Vivo. Biology Open 2013, 2 (3), 306–313. https://doi.org/10.1242/bio.20133665.

(36) Nin, V.; Hernández, J. A.; Chifflet, S. Hyperpolarization of the Plasma Membrane Potential Provokes Reorganization of the Actin Cytoskeleton and Increases the Stability of Adherens Junctions in Bovine Corneal Endothelial Cells in Culture. Cell Motility 2009, 66 (12), 1087–1099. https://doi.org/10.1002/cm.20416.

(37) Chifflet, S.; Hernández, J. A.; Grasso, S.; Cirillo, A. Nonspecific Depolarization of the Plasma Membrane Potential Induces Cytoskeletal Modifications of Bovine Corneal Endothelial Cells in Culture. Experimental Cell Research 2003, 282 (1), 1–13. https://doi.org/10.1006/excr.2002.5664.

(38) Sharma, M.; Astekar, M.; Soi, S.; Manjunatha, B. S.; Shetty, D. C.; Radhakrishnan, R. PH Gradient Reversal: An Emerging Hallmark of Cancers. Recent Pat Anticancer Drug Discov 2015, 10 (3), 244–258. https://doi.org/10.2174/1574892810666150708110608.

