## Supplemental Figures for "An optogenetic tool to raise intracellular pH in single cells and drive localized membrane dynamics"

**Author List:** Caitlin E. T. Donahue[a,b], Michael D. Siroky[a,b], and Katharine A. White[a,b*]

**Institution Address:**

[a]Department of Chemistry and Biochemistry, University of Notre Dame, Notre Dame, IN 46556, United States

[b]Harper Cancer Research Institute, University of Notre Dame, South Bend, IN 46617, United States

**Corresponding author:**

*

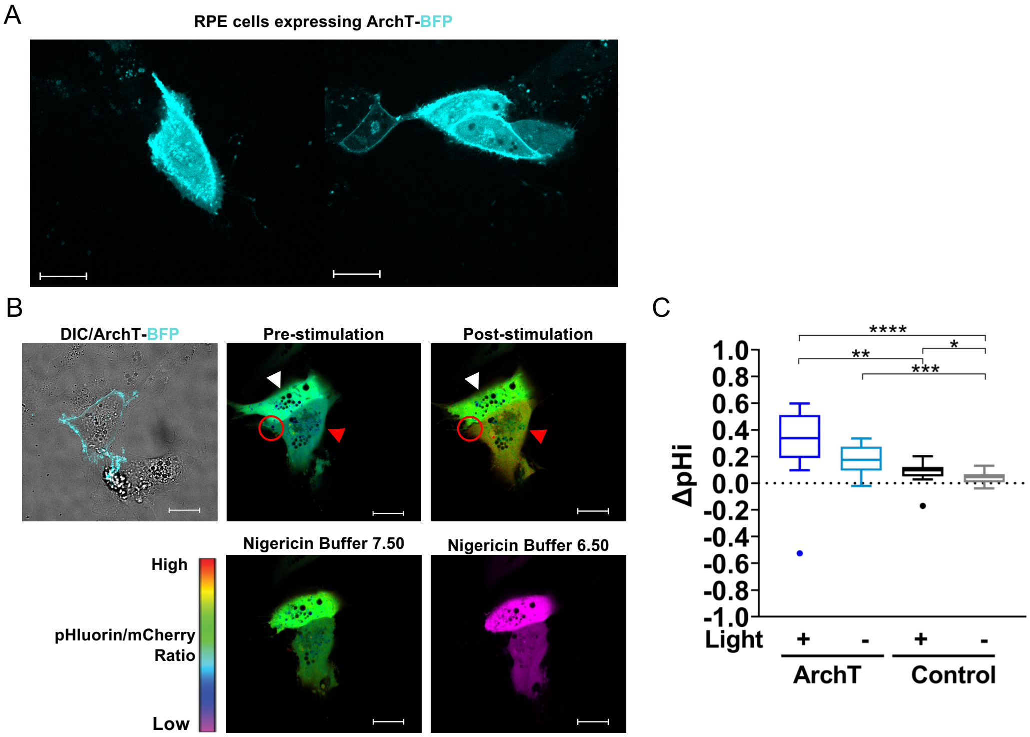


**Figure S1. Quantification of Archaerhodopsin pHi increases using ratiometric mCherry-pHluorin imaging. (A)** Representative images of RPE cells expressing ArchT-BFP. Scale bar 20 µm. **(B)** An ArchT RPE cell expressing mCherry-pHluorin (red arrow) and control RPE cell expressing mCherry-pHluorin (white arrow) are both photoactivated with 561 nm light within the stimulation region of interest (ROI, red circle). Ratiometric display of pHluorin/mCherry fluorescence ratios pre-stimulation, post-stimulation, and during pH standardization (see methods for details). Scale bar 20 µm. **(C)** Quantification of pHi changes for stimulated (+Light) and unstimulated (-Light) cells prepared as described in (B). Tukey boxplots (n = 15-55 cells per condition, 2 biological replicates), significance determined using the Kruskal-Wallis test, Dunn's multiple comparison correction. * $p<0.05$, ** $p<0.01$, *** $p<0.001$, **** $p<0.0001$.

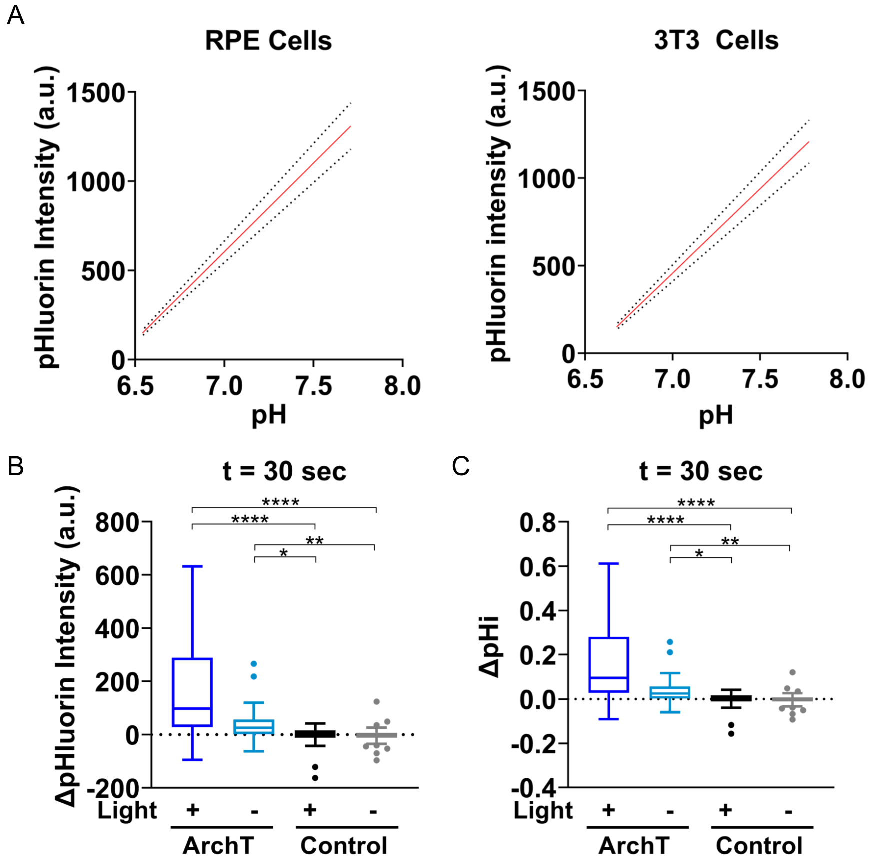


**Figure S2. Standard curves for back-calculation of pHi data from pHluorin intensity. (A)** Standard curve data from pHluorin intensity measurements in RPE and NIH-3T3 cells. The red line indicates the mean while the dotted black lines represent the 95% confidence interval. (n=261 cells for RPE, n=64 cells for 3T3, 2 biological replicates). **(B)** Additional quantification from Figure 1E. Quantification of pHluorin intensity changes at 30 seconds into experiment (30% LED power). **(C)** Quantification of pHi changes in cells in (B) using standard curves in (A). For (B-C) Tukey boxplots (n= 26-62, 3-5 biological replicates). Significance determined using the Kruskal-Wallis test, Dunn's multiple comparison correction, *p<0.05, **p<0.01, and ****p<0.0001

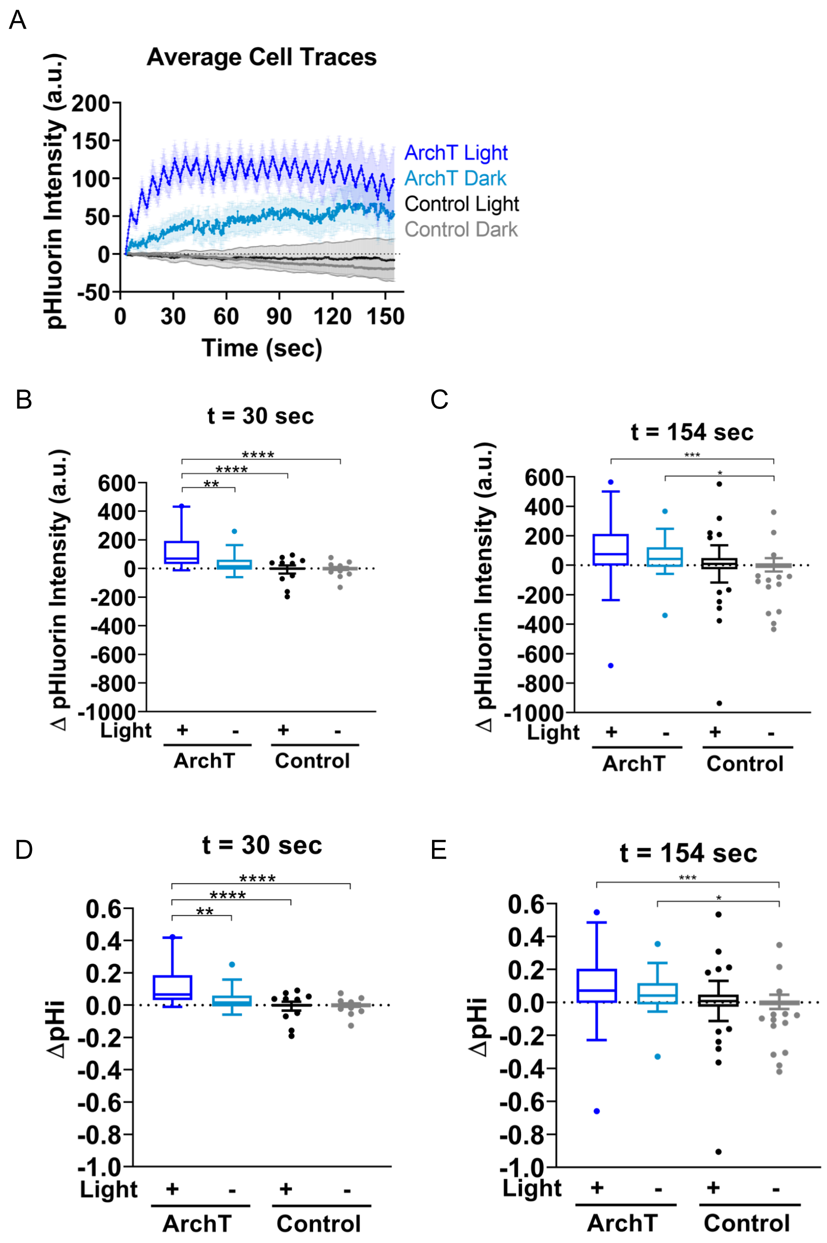


**Figure S3: Archaerhodopsin can spatiotemporally increase pHi in single RPE cells using 100% LED Power. (A)** Average pHluorin intensity traces over the course of the 100% LED power stimulation experiment. Traces show mean ± SEM (n = 20-69 cells from 3-5 biological replicates). **(B-C)** Quantification of pHluorin intensity changes at 30 sec **(B)** and the end **(C)** of the experiment in (A). **(D-E)** Quantification of pHi changes at 30 sec **(D)** and the end **(E)** of the experiment in (A). For (B-E), Tukey boxplots shown with significance determined using the Kruskal-Wallis test, Dunn's multiple comparison correction, *p<0.05, **p<0.01, ***p<0.001, and ****p<0.0001.

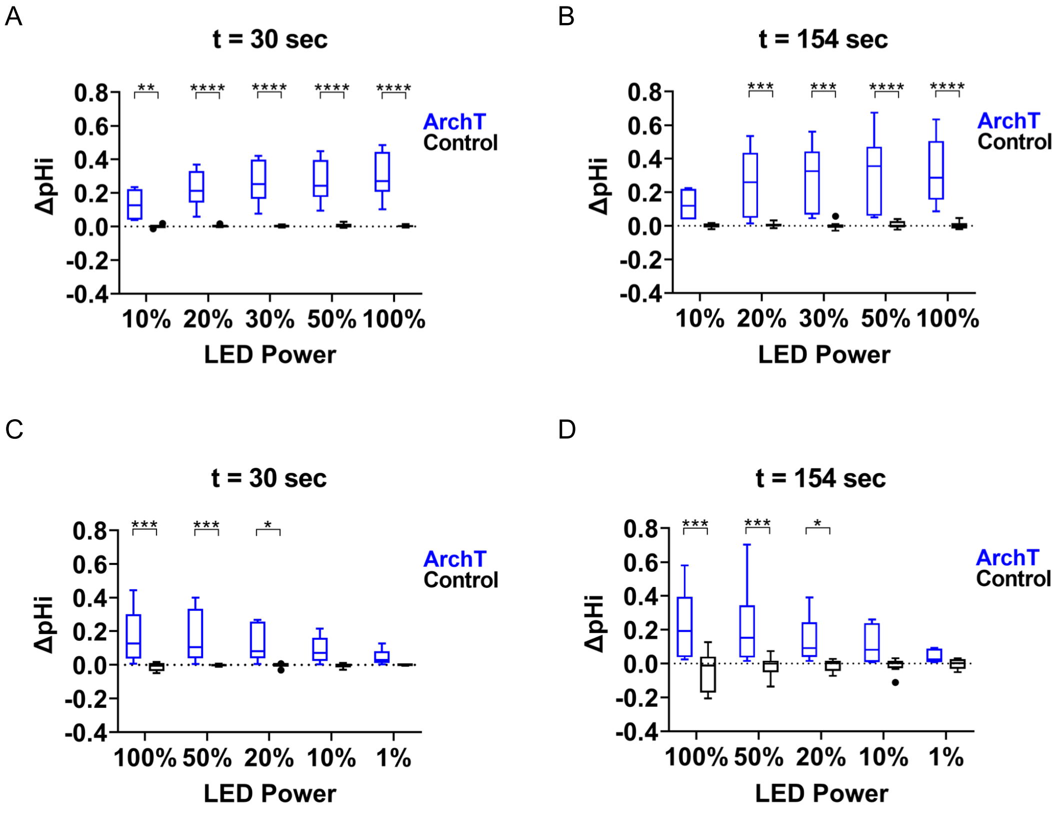


**Figure S4: ArchT increases pHi at a range of LED stimulation powers**. Quantification of pHi change using an increasing sequence (10%, 20%, 30%, 50%, 100%) of LED stimulation power titration at **(A)** 30 seconds and **(B)** at the end of the experiment (154 seconds). Quantification of pHi using a decreasing LED stimulation power sequence (100%, 50%, 20%, 10%, 1%) at **(C)** 30 seconds and **(D)** at the end of the experiment (154 seconds). For (A-D) Tukey boxplots (n= 6-8 cells per condition from 1-3 biological replicates), significance determined using two-way ANOVA and Holm-Šídák's multiple comparison test. *p<0.05, **p<0.01, ***p<0.001, and ****p<0.0001.

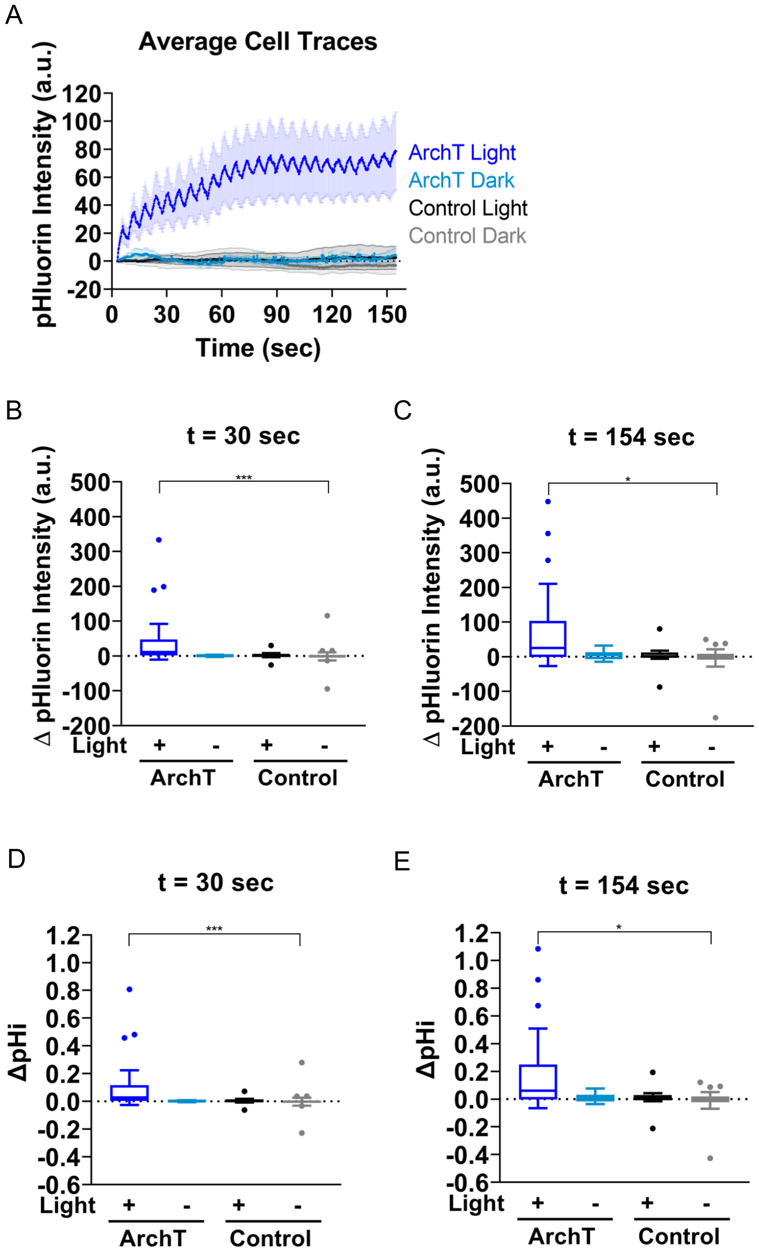


**Figure S5: Archaerhodopsin can spatiotemporally increase pHi in single NIH-3T3 cells. (A)** Average pHluorin intensity traces over the course of the stimulation experiment. Traces show mean ± SEM (n = 9-31 cells per condition from 3 biological replicates). **(B-C)** Quantification of pHluorin intensity changes at 30 sec **(B)** and the end **(C)** of the experiment in (A). **(D-E)** Change in pHi quantified for cells at 30 sec **(D)** and at the end **(E)** of the experiment in (A). For B-E, Tukey boxplots, n = 9-31 per condition significance determined using the Kruskal-Wallis test, Dunn's multiple comparison correction, *p<0.05, and ****p<0.0001.

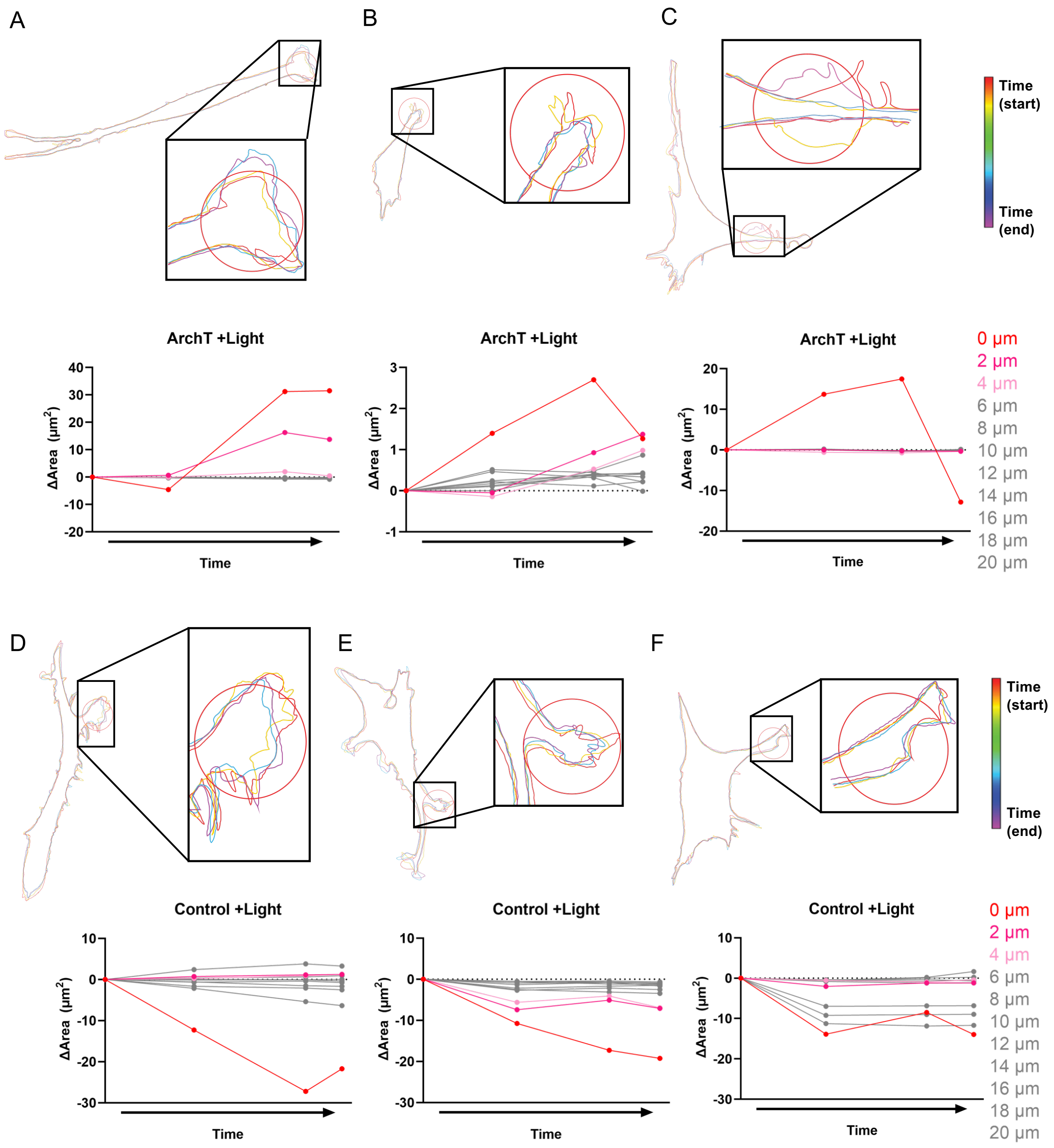


**Figure S6. Additional Membrane Responses for NIH-3T3 cells treated as in Figure 3.** Additional representative images of ArchT **(A-C)** and Control expressing **(D-F)** NIH-3T3 cells stimulated with 561 nm light. As in Figure 3, membranes were traced at key movie frames and overlaid, t= 0 sec (red) to 154 sec (violet). Zoomed-in sections of the stimulation region and the membrane dynamics are also provided. As in Figure 4, change in area for segmented cells are located below their respective trace. 0 µm indicates the stimulation ROI with each subsequent segment representing 2µm intervals.

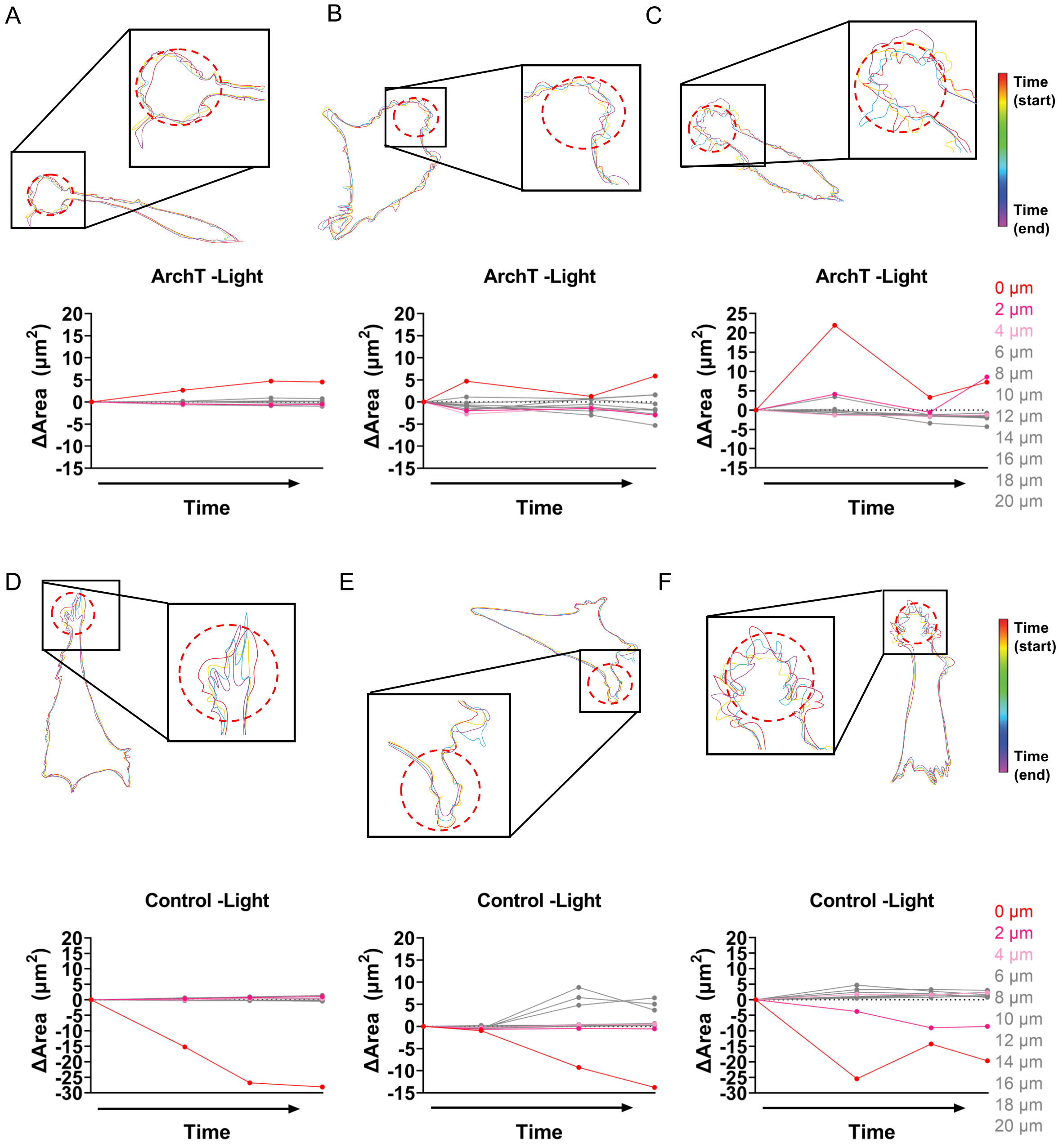


**Figure S7. Additional Membrane Responses for NIH-3T3 cells mock stimulated as in Figure 3.** Additional representative images of ArchT **(A-C)** and Control expressing **(D-F)** NIH-3T3 cells mock-stimulated (no 561 nm light). As in Figure 3, membranes were traced at key movie frames and overlaid, t= 0 sec (red) to 154 sec (violet). Zoomed-in sections of the stimulation region and the membrane dynamics are also provided. As in Figure 4, change in area for segmented cells are located below their respective trace. 0 μm indicates the stimulation ROI with each subsequent segment representing 2μm intervals.

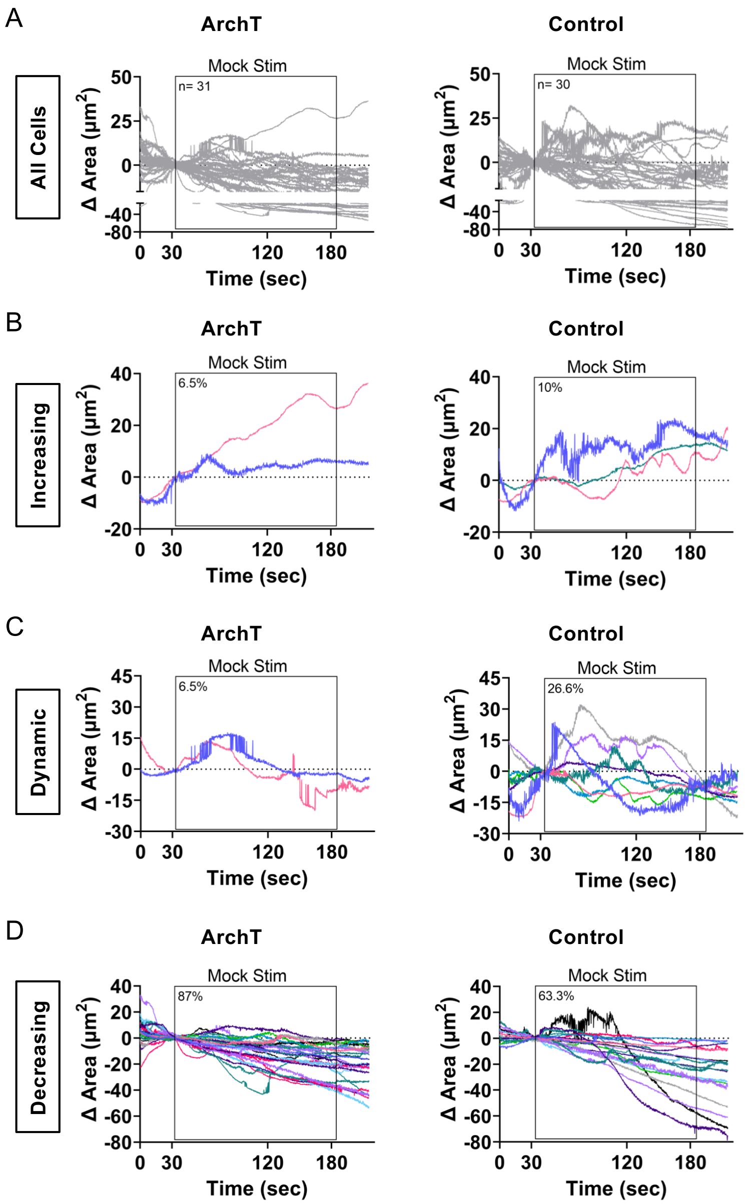


**Figure S8. Membrane dynamics with a mock stimulation. (A)** Membrane area changes were quantified within a mock-stimulation ROI for ArchT and control NIH3T3 cells. Individual traces are shown for each cell, black box indicates timing of mock stimulation (n=30-31 cells per condition, from 3 biological replicates). **(B-D)** Binned traces of mock-stimulated ArchT and control 3T3 cells in (A) to characterize phenotypes that are protrusive **(B)**, dynamic **(C)**, and decreasing/static **(D)**. Percentage of cells binned in each category is shown on each graph and summarized in Figure 5E.
